# Torch-eCpG: A fast and scalable eQTM mapper for thousands of molecular phenotypes with graphical processing units

**DOI:** 10.1101/2023.03.07.531597

**Authors:** Kord M. Kober, Liam Berger, Ritu Roy, Adam Olshen

## Abstract

**Background:** Gene expression may be regulated by the DNA methylation of regulatory elements in *cis, distal*, and *trans* regions. One method to evaluate the relationship between DNA methylation and gene expression is the mapping of expression quantitative trait methylation (eQTM) loci (also called expression associated CpG loci, eCpG). However, no open-source tools are available to provide eQTM mapping. In addition, eQTM mapping can involve a large number of comparisons which may prevent the analyses due to limitations of computational resources. Here, we describe Torch-eCpG, an open-source tool to perform eQTM mapping that includes an optimized implementation that can use the graphical processing unit (GPU) to reduce runtime.

**Results:** We demonstrate the analyses using the tool are reproducible, up to 18x faster using the GPU, and scale linearly with increasing methylation loci.

**Conclusions:** Torch-eCpG is a fast, reliable, and scalable tool to perform eQTM mapping. Source code for Torch-eCpG is available at https://github.com/kordk/torch-ecpg.

## BACKGROUND

Gene expression is regulated, in part, by epigenetic mechanisms. A major unanswered question in genomics research is the functional contribution of epigenetic variation on gene expression.[1, 2] One method to evaluate for the potential functional effect of a methylation variation is to test for an association between levels of methylation and gene expression from the same samples. These expression-associated quantitative trait methylation (eQTM) loci may contribute to the regulation of gene expression (also called expression associated CpG loci, eCpG). These associations may be local (e.g., methylation located in the promoter region of a gene) or remote (e.g. methylation loci in a distant enhancer regions of a gene or on a different chromosome). There is growing interest in the integration of these data modalities and evaluating for eQTMs. For example, in terms of clinical research, recent studies have identified eQTMs from a variety of tissue types and outcomes.[3-7]

Recent advances in high throughput molecular methods allow for the collection of complementary methylation and gene expression data from the same sample in large numbers. Although an increasing number of recent studies have provided eQTM datasets,[3, 8] there are no open-source tools currently available to investigators to implement these analyses on their own.

Given the increase in the availability of complementary datasets and the biological utility of identifying eQTMs, analytic tools must be made available, free to use, and able to scale to handle thousands to millions of samples (e.g., patients or single cells). Current methylation array datasets provide hundreds of thousands of loci (e.g., Infinium MethylationEPIC, Illumina, San Diego, CA) and RNA-sequencing and microarray methods provide expression levels for tens of thousands of genes. An exhaustive evaluation of these datasets would result in tens of billions of tests for hundreds or thousands of samples, quickly outreaching the computing capacity of a desktop computer and requiring larger workstations, clusters, or cloud computing.[9] Future datasets will likely include more loci for evaluation and larger sample sizes. This need for computational resources will also require improvements in efficiency. Graphical processing units (GPUs) have provided major improvements in computation efficiency (i.e., runtime) for many bioinformatic software tools.[10] Readily available open source libraries implement numerous general-purpose methods and mathematical primitives that allow for major improvements in computational efficiency at relatively lower costs as compared to CPUs.[11]

Given the analytic utility of evaluating for eQTMs to identify relationships between gene expression and epigenetic changes, a lack of an available open-source tool to implement an eQTM analysis, and the performance benefits of utilizing a GPU, the objectives of this project were to develop an open source, general-use tool for eQTM mapping and evaluate for performance increases of a GPU implementation. Here, we present the Torch-eCpG tool (tecpg).

## IMPLEMENTATION

### Association analyses

Two methods are available to test for the associations between CpG methylation and gene expression (eCpGs). First, a Pearson correlation can be computed between the methylation level and gene expression level. Second, a multivariate linear regression (MLR) method can model the relationship between gene expression and methylation level while including while adjusting for covariates (e.g., age, batch).[4, 8, 12] We tested for an association between methylation at CpG *j* and the expression level of transcript *k*, by fitting the model

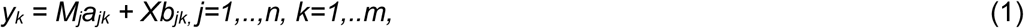

where *y*_*k*_ is a vector of log expression levels at gene *k* with length *m, M*_*j*_ is a size *n* vector of methylation values (i.e., Beta scores) at CpG *j, m* is the number of covariates, and *X* is a *n* x *m* matrix of covariates.

Given the PyTorch toolkit does not currently include a function to estimate the cumulative distribution function (CDF) for the Student’s t-distribution, and thus it is not easily possible to compute a p-value based on the t-distribution, we used a Gaussian distribution CDF to estimate p-values. The MLR feature of tecpg is optimized to increase performance for large input datasets. Optimizations include the minimization of repeated calculations, parallelizing tasks, memory use management, and selective use of tensors on the GPU.

Gene expression, methylation, and phenotypic (i.e., covariate) data are provided as comma-separated value (CSV) files. Gene and methylation loci genomic region annotations are provided as browser extensible data (BED) files. Four eCpG mappings modalities are implemented:

- *Cis*-eCPG: associations between all methylation loci-gene expression pairs within a specified window (default + 1 Mb) around the transcript start site for genes.
- *Distal*-eCpG: associations between all methylation loci-gene expression pairs outside of a specified window (default 50 Kb) from the transcript start site for genes, but on the same chromosome.
- *Trans*-eCpG: associations between all methylation loci-gene expression pairs. The computation is performed for each chromosome using methylation loci on all other chromosomes. To reduce output size, only associations below a given *p* value threshold (default 1x^10-5^) are stored.
- *All*-*by-all*: associations between all methylation loci-gene expression pairs across all regions. To reduce output size, only associations below a given *p* value threshold (default 1x^10-5^) are stored.

Numerous user-friendly features are provided. The tool will attempt to automatically detect a CUDA supported GPU. If a supported GPU is not available, or upon user request, the analyses will be performed using a CPU. The number of CPU threads is configurable and threaded CPU processing is available. In the case where data sizes exceed the CPU or GPU memory, the tool can be set to batch the analyses into chunks of gene expression and/or methylation data. An option is available to estimate the number of gene expression loci per chunk. Finally, to limit the size of the output and associated time writing the file out, the user can set a p-value threshold to filter the reported analyses and can select the columns of the MLR analyses to report.

### Evaluation

To evaluate for the replicability of the regression analyses implemented in tecpg, we compared our regression analyses with similar analyses using the cor() and lm() functions in the stat package in R. To benchmark tecpg, we compared *cis*-eCpG, *distal*-eCpG and *trans*-eCpG mapping performance with and without a GPU. For CPU-based comparisons of the individual regions, computations were limited to a single core.[11] We also evaluated tecpg performance using a range of CPU core counts (i.e., 1, 2, 4, 8, 16, 24). These analyses used a dataset of whole blood samples collected from 333 participants (76% female) aged 18–78 years in the Grady Trauma Project (GTP) (Gene Expression Omnibus (GEO) accession numbers GSE72680, GSE58137). To facilitate the evaluation of the scaling performance of the GPU implementation when mapping *trans*-eCpGs across a wide range of eCpG counts, we sampled with replacement from the GTP dataset to obtain a sample size of 1000. All tecpg benchmarks were conducted on a physical server running Linux having 28 Xeon cores (2.3GHz), 256 GB CPU memory, and a A2 GPU with 16GB of memory (Nvidia Corporation, Santa Clara, CA).

## RESULTS AND DISCUSSION

To provide an open-source tool for eQTM mapping, we developed the Torch-eCpG software package. To evaluate the reproducibility of the linear regression analysis, we compared our results with those implemented in the lm() function in the stats package in R. As shown in Figure 1, our implementation of the linear regression demonstrates high reproducibility.

**Figure 1.**
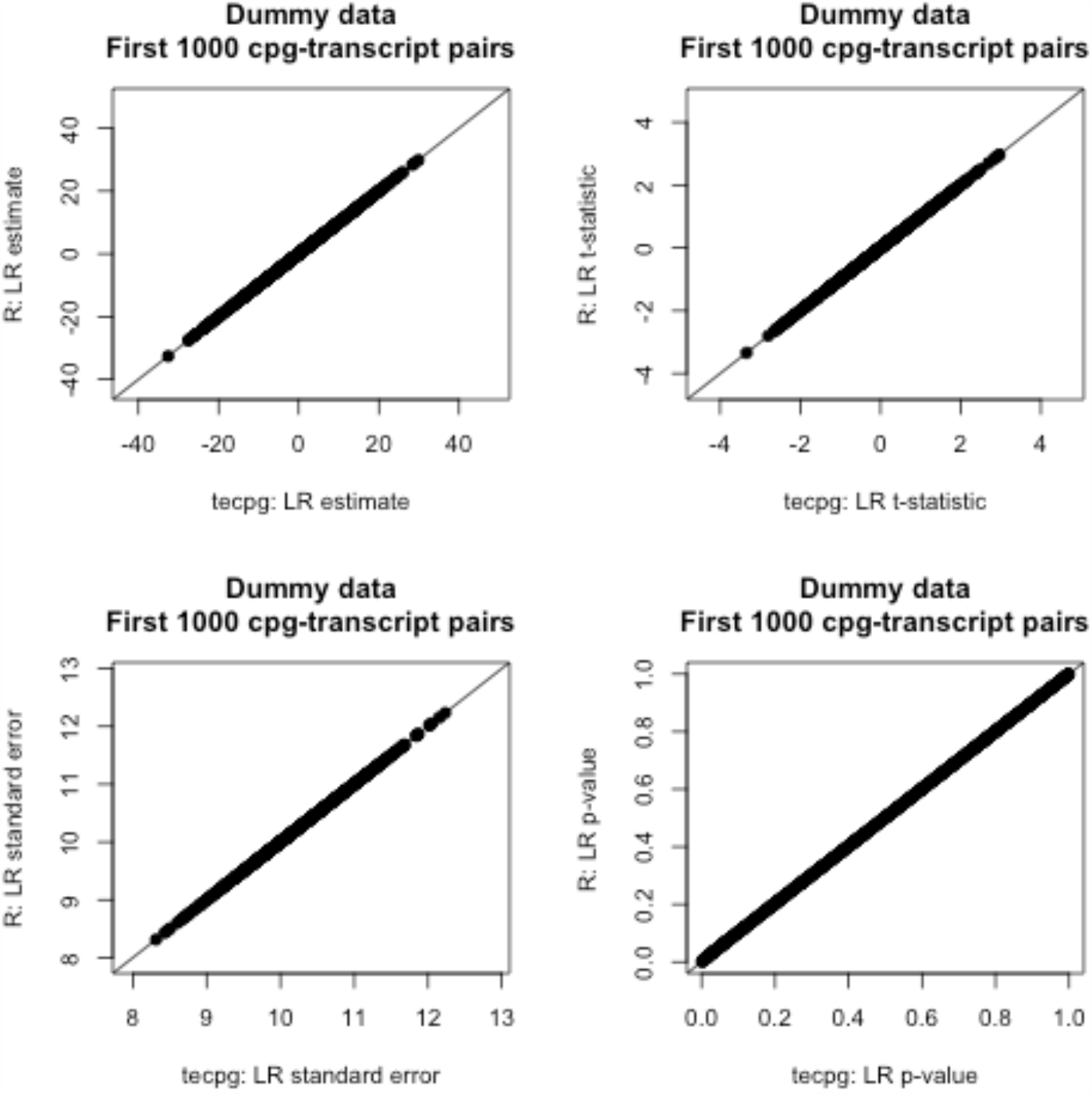
Comparisons of the first 1000 CpG-transcript pair linear regression analyses results between tecpg and lm() function in the stats package in R for a simulated dataset generated by sampling with replacement (n=1000 samples). Simulated patient data was generated from real patient data in the Grady Trauma Project.

For tecpg benchmarking, we evaluated eQTM mapping using the CPU and GPU implementation in tecgp for 300 patients from the GTP patient dataset (422,442 methylation loci and 17,653 genes). Across the mapping modalities, the GPU outperformed the CPU analysis by up to 18x. Our implementation of the *cis-*eCpG mapping was 1.4x faster on the GPU than that of the CPU (Figure 2A). For *distal-*eCpG mapping, our implementation was 5x faster on the GPU. Finally, for *trans*-eCpG mapping, our implementation was 18x faster on the GPU. In terms of tecpg using additional cores, we found that major incremental improvements were realized by increasing the CPU core count up to 8, after which the gains were minimal (Figure 3). Although the CPU performance did improve with additional cores, the GPU implementation was still 2x faster than the 24-core CPU implementation.

**Figure 2.**
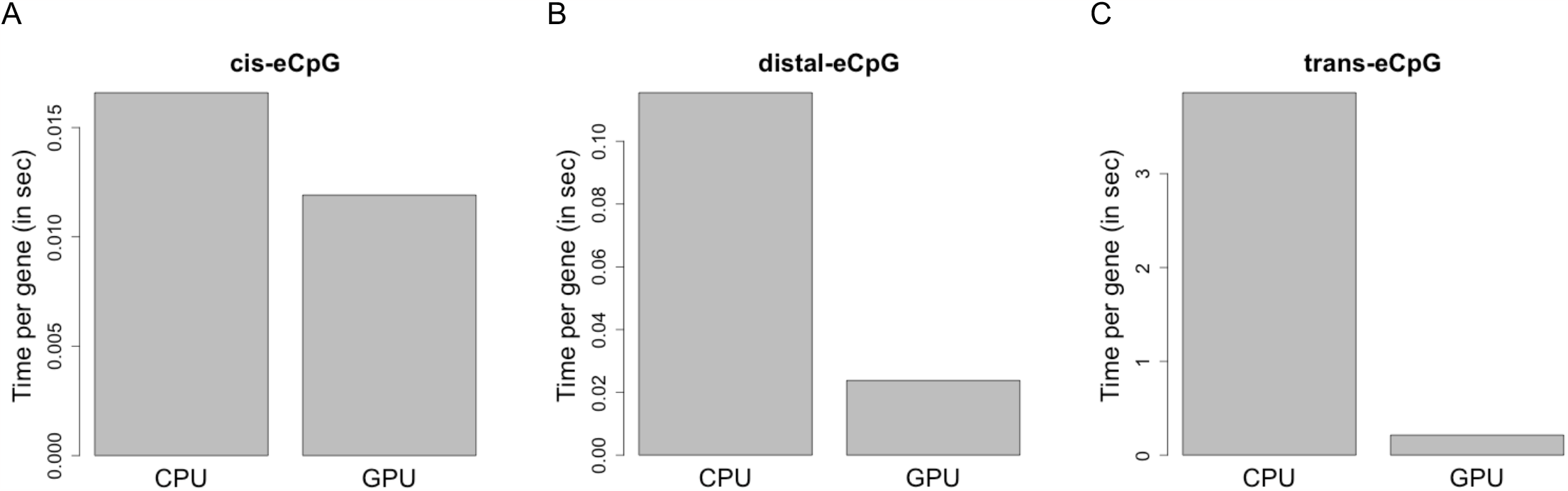
Performance of GPU implementations for eQTM mapping. Comparison of runtimes for tecpg analyses on CPU and GPU for (A) *cis*-eCpG, (B) *distal*-eCpG, and (C) *trans*-eCpG. The analyses evaluated 340 patients from the Grady Trauma Project dataset and included 422,442 methylation loci and 17,653 genes.

**Figure 3.**
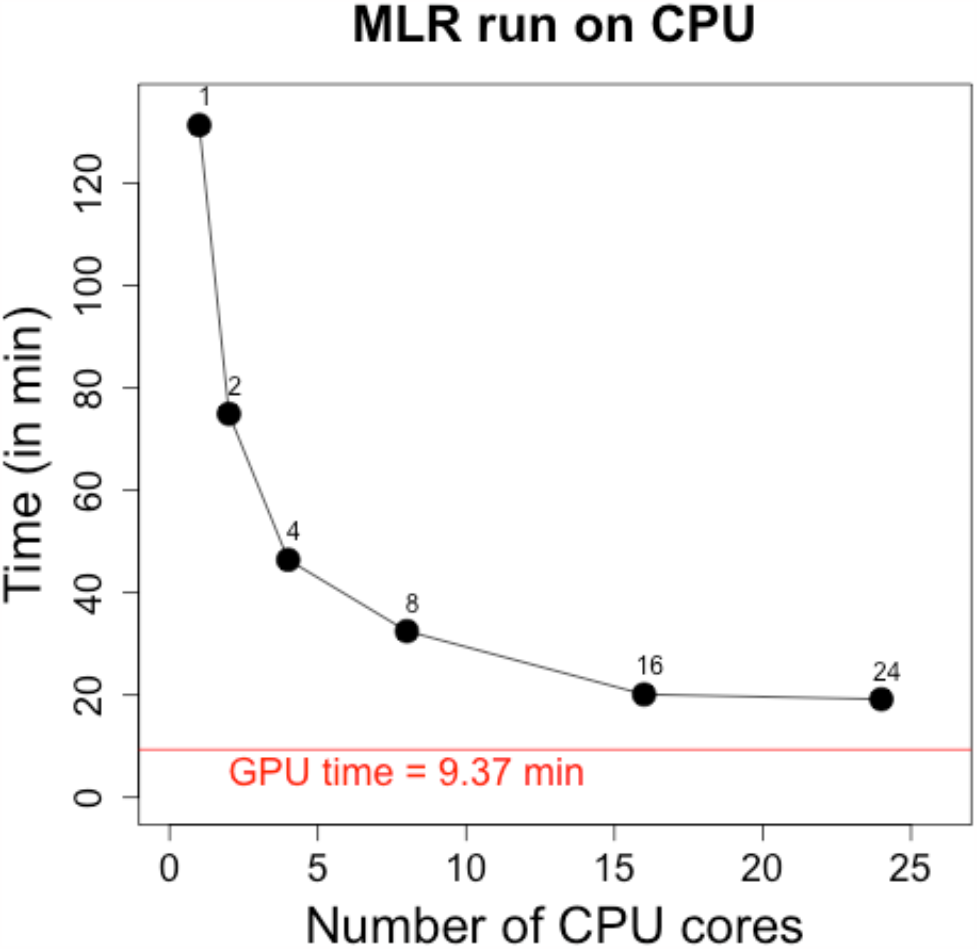
CPU runtimes for tecpg using 1, 2, 4, 8, 16, and 24 CPU cores. The analyses evaluated 340 patients from the Grady Trauma Project dataset and included 422,442 methylation loci and 17,653 genes.

We found tecpg scales linearly across a wide range of methylation loci for a reasonably large sample size (n=1000 patients) (Figure 4). In addition, the total time to evaluate 1000 patients for whole transcriptome (2×10^4^ genes) and whole methylome array data (8.5×10^5^) was <15 hours. The short time needed to evaluate a dataset sized to the largest currently available methylation array (i.e., Infinium MethylationEPIC) highlights the utility of this tool to evaluate eQTM mapping of dataset of realistic size. The linear scaling suggests the tool will be useful for larger datasets in the future (e.g., >10,000 patients).

**Figure 4.**
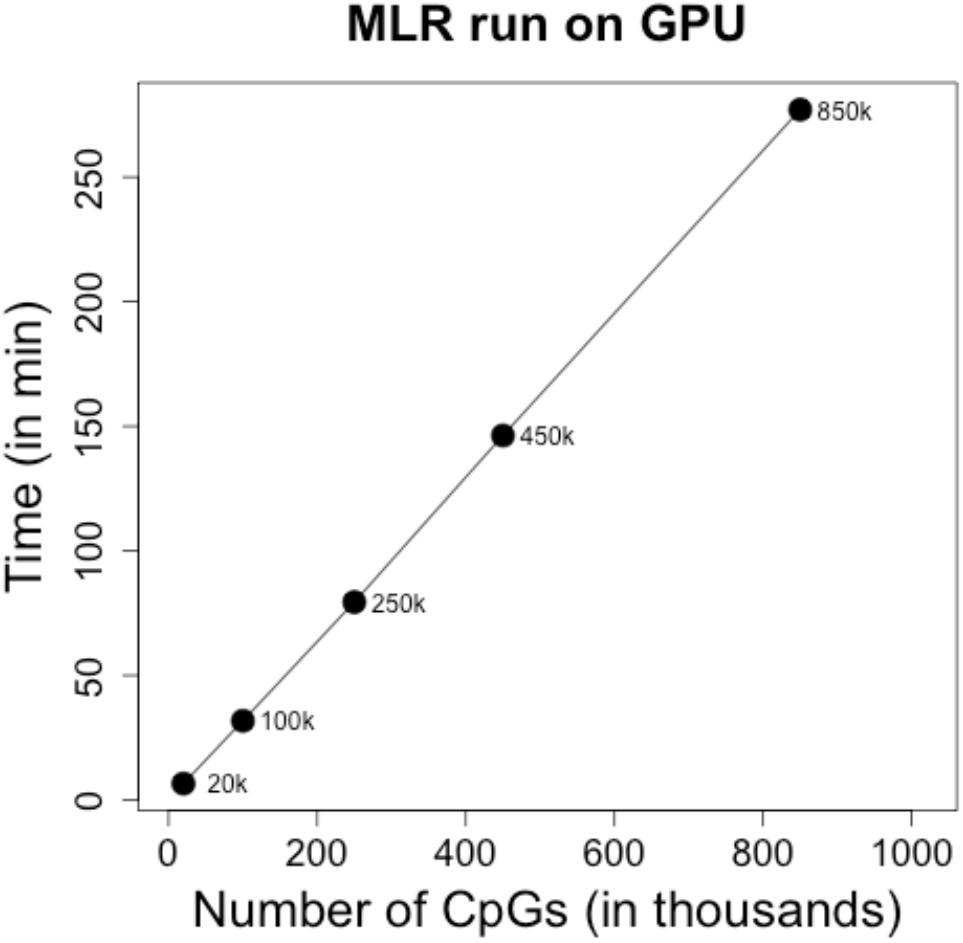
GPU runtime of tecpg for 1000 simulated patient samples for 20,000 genes and 20×10^3^, 100 ×10^3^, 250 ×10^3^, 450 ×10^3^ and 850 ×10^3^ CpG loci. Simulated patient data was generated from real patient data in the Grady Trauma Project.

## CONCLUSIONS

Torch-eCpG is the first freely available open-source tool to perform eQTM mapping. It provides a scalable and high-performance implementation that supports GPU enabled systems. By reducing computing time the tool offers cost-savings on shared systems (e.g., clusters) or cloud-based computing resources that charge by units of time. This tool allows for individual research labs with limited computational resources to perform analyses on affordable computer equipment or cloud-based virtual machines.

## AVAILABILITY AND REQUIREMENTS

Project name: Torch-eCpG

Project home page: http://www.github.com/kordk/torch-ecpg

Operating system(s): Platform independent

Programming language: Python 3.10 or higher

Other requirements: click∼=8.0.3, colorama∼=0.4.4, matplotlib∼=3.5.1, numpy∼=1.24.1, pandas∼=1.3.5, psutil∼=5.9.4, requests∼=2.26.0, scipy∼=1.10.0, setuptools∼=63.3.0, torch∼=1.13.1+cu116

License: BSD-3-Clause

Any restrictions to use by non-academics: license needed

## LIST OF ABBREVIATIONS

BED: Browser extensible data
CDF: Cumulative distribution function Cp
G: Cytosine phosphate guanine
CPU: Central processing unit
CSV: Comma separated value
DNA: Deoxyribonucleic acid
eCpG: Expression associated CpG
eQTM: Expression quantitative trait methylation
GEO: Gene Expression Omnibus
GB: Gigabyte
GHz: Gigahertz
GPU: Graphical processing unit
GTP: Grady Trauma Project

## DECLARATIONS

### Ethics approval and consent to participate

Not applicable.

### Consent for publication

Not applicable.

### Availability of data and materials

The datasets analyzed in this study are publicly available in the Gene Expression Omnibus repository (https://www.ncbi.nlm.nih.gov/geo/query/acc.cgi?acc=GSE72680 and https://www.ncbi.nlm.nih.gov/geo/query/acc.cgi?acc=GSE58137)

### Authors’ contributions

KMK originated the idea and developed the initial project plan. KMK and EB outlined and planned the tool development. EB and KMK developed the Torch-eCpG tool. All authors were involved in the planning and testing of Torch-eCpG. KMK prepared and reviewed the manuscript and figures. All authors reviewed and edited the manuscript.

## Acknowledgements

Not applicable.

## Authors’ information (optional)

Not applicable.

